# A Highly Protective Clade 1 and 2 Cross-Reactive Pandemic Influenza Virus Vaccine Based on a 4th Generation Fully Deleted Adenoviral Vector of a Rare Serotype

**DOI:** 10.1101/2025.04.30.651579

**Authors:** Yan Qi, Richard Bowen, Airn Tolnay Hartwig, Janae Wheeler Cull, Jessica Kelly, Xenia Klimenova, Uwe D Staerz

## Abstract

The GreVac vaccine technology was created as a fast and flexible plug-and-play vaccine platform based on a 4th generation architecture of fully deleted (fd) helper virus-independent (hi) adenoviral (Ad) vectors. For the initial proof-of-principle studies, we at Greffex had engineered an avian influenza vaccine, which delivered a transgene expression cassette for an avian influenza virus H5 hemagglutinin and N1 neuraminidase genes in a capsid of the common human Ad serotype 5 (Ad5). This vaccine proved highly immunogenic and protective in mice. These studies revealed that *intramuscular* (*i.m.*) delivery proved more efficient than *subcutaneous* (*s.c.*) or *intranasal* (*i.n.*) routes. In the human population, pre-exposure to the Ad5 virus is common. To minimize interference by pre-existing anti-Ad5 immunities, we created a new GreVac-based avian influenza vaccine, in which the fd Ad genome was packaged into a capsid of the rare human Ad serotype 6 (Ad6). We now report that at very low doses, the resulting GreFluVie6 vaccine given *i.m.* fully protected mice and ferrets against lethal challenges with the clade 1 *A/Vietnam/1203/2004* avian influenza virus associated with induction of potent immune cellular and humoral immune responses. The recipients’ serum antibodies strongly cross-reacted with clade 2.1.3.2 (*A/Indonesia/05/2005*) and clade 2.3.4.4b H5 hemagglutinins.

## BACKGROUND

Influenza is a group of infectious diseases of the respiratory tract caused by RNA viruses of the family Orthomyxoviridae and genera A, B and C (Kamps et al., 2006; Krammer et al., 2018). The natural reservoir of influenza A viruses can be found in wild birds (Venkatesh et al., 2018). The structure of the influenza virus genome expedites the appearance of virus variants through genetic shifts and drifts (Dadonaite et al., 2019; Kamps et al., 2006; Krammer et al., 2018; Olsen et al., 2006; Parrish et al., 2015). Every 10 to 50 years, highly pathogenic influenza strains have emerged, some of which have caused millions of deaths (Kamps et al., 2006). The highly lethal H5 clade 1 *A/Vietnam/1203/2004* avian influenza emerged in Asia in 2003 (He & Kam, 2024; Kamps et al., 2006; Sutton, 2018). Fortunately, it never acquired an ability for efficient human-to-human transmission. Nevertheless, one H5 clade 2.1 variant, *A/Indonesia/05/2005*, spread throughout large regions of the world with Egypt being its hotspot (Refaey et al., 2015). Recently, clade 2.3.4.4b H5 influenza virus variants appeared that possess a broad infectivity to numerous different animal species (Kandeil et al., 2023; Webby & Uyeki, 2024). This virus has been spreading throughout the world, now threatening domestic livestock such as cows and chickens (Graziosi et al., 2024). To date, human infections with the 2.3.4.4b virus have occurred through direct contact with infected animals and sustained human-to-human transmission has not been reported (Ueki et al., 2025). However, several gain-of-function studies with the *A/Vietnam/1203/20*04 avian influenza virus demonstrated that that only few mutations were required to expand its infectivity (Herfst et al., 2012; Imai et al., 2012; Russell et al., 1977). Therefore, a more widespread human influenza pandemic with a clade 2.3.4.4b H5 influenza virus may indeed become reality once the underlying virus had acquired efficient human transmission. (Ison & Marrazzo, 2025).

Highly infectious influenza viruses can spread through a human population faster than present technologies deliver new vaccines (Buckland, 2015; Lambert & Fauci, 2010). We decided to optimize the fully deleted Ad technology into a fast and flexible plug-and-play vaccine platform. The resulting GreVac vectors are based on a proprietary architecture of fully deleted (fd) helper virus-independent (hi) adenoviral (Ad) constructs of different serotypes (Lee et al., 2019; Qi, Zhang, et al., 2024). Their genomes are deleted of all endogenous Ad genes to curtail the induction of and interference by anti-Ad responses. Removing the StaerzAd genes better focuses the immune system to specific antigens (Abbink et al., 2016; Weaver et al., 2009, 2013), increases the vaccine efficacy (Qi et al., 2024), and enables efficient short-term prime-boost vaccination protocols (Koehler et al., 2006; Qi et al., 2024). For the initial proof-of-principle studies of the GreVac approach, an avian influenza vaccine was engineered. It delivered a transgene expression cassette for the avian influenza virus H5 hemagglutinin and N1 neuraminidase genes in a *gutted* Ad genome that was packaged in a capsid of the common human Ad serotype 5 (Ad5) (Qi, Tarbet, et al., 2024). This vaccine proved highly immunogenic and protective in mice against challenges with the clade 1 *A/Vietnam/1203/2004* avian influenza virus. It demonstrated its highest efficiency when given via an *i.m.* but not *s.c. i.n.* administraton. Once we had demonstrated that GreVac-based vaccines were highly active, we further optimized their designs for human applications. In humans, infections with viruses of the Ad5 serotype are quite common (Akello et al., 2020) and elicit immune responses that hamper the activity of subsequently delivered Ad5-based vectors (Fausther-Bovendo & Kobinger, 2014). Therefore, we chose an Ad capsid of the rare human serotype 6 (Ad6) to deliver the fully deleted Ad genome (Akello et al., 2020). As previously discussed in detail (Qi, Zhang, et al., 2024), the GreVac system entails packaging of the fully deleted Ad genome and is not mediated by helper virus constructs (Cheshenko et al., 2001; Palmer & NG, 2005; Palmer & Ng, 2003). This approach is based on two independently modifiable components. They are a fully gutted GreVac vector genome and a non-packageable circular expression plasmid. Vector production is initiated by the co-transfection of both components into engineered host cells (Qi et al., 2024). Our system limits the frequent recombination of replication competent adenoviruses (RCA) or the presence of helper viruses, both of which were seen with earlier iterations of the fd Ad technology (Cheshenko et al., 2001; Dormond et al., 2009, 2010). These contaminants are by themselves immunologically active and may inhibit the vector function (Nomura et al., 2004). In the avian influenza vaccine of this proceeding, GreFluVie6, the genome carries a transgene expression cassette of the clade 1 *A/Vietnam/1203/2004* avian influenza virus H5 hemagglutinin and N1 neuraminidase genes. It is packaged in a capsid of the rare human Ad serotype 6 (Ad6) to minimize interference by pre-existing immune responses of the human recipients. The resulting GreFluVie6 vaccine given *i.m.* fully protected mice and ferrets against lethal challenges with *A/Vietnam/1203/2004* at low doses. It induced potent humoral and cellular immune responses against the vaccine antigens. We also found that the recipients’ serum antibodies strongly cross-reacted with clade 2.1.3.2 (*A/Indonesia/05/2005*) and clade 2.3.4.4b H5 hemagglutinins.

## RESULTS

### Composition of the GreFluVie6 Vaccine

The GreVac-based fd genome of GreFluVie6 was previously described in detail (Qi et al., 2024). It contains a bicistronic expression cassette for the hemagglutinin 5 (H5) (AY818135.1) and neuraminidase 1 (N1) (AY651447.1) of the clade 1 *A/Vietnam/1203/2004* influenza virus. It is flanked by inverted terminal repeats (ITR) and a packaging signal Y both derived from an Ad of the human serotype 5. To increase the genome to a packageable size, a DNA stuffer was added that was derived from internal sequences of the human house-keeping gene 5-aminoimidazole-4-carboxamide ribonucleotide formyl-transferase/IMP cyclohydrolase (ATIC). Once released from the cloning plasmid by a restriction enzyme cut, the linear GreFluVie genome was packaged into an Ad capsid of the human serotype 6 with the help of the packaging expression plasmid pPaC6 (**Figure 1**). The linear genome and a packaging expression plasmid were co-transfected into a variant of the human embryonic kidney (HEK293) cells (Qi, Tarbet, et al., 2024). The completed GreFluVie6 vaccine vector was released from the production cells and purified by a modification of a two-step column chromatography into a vector formulation buffer (VFB) (Eglon et al., 2009).

**Figure 1.**
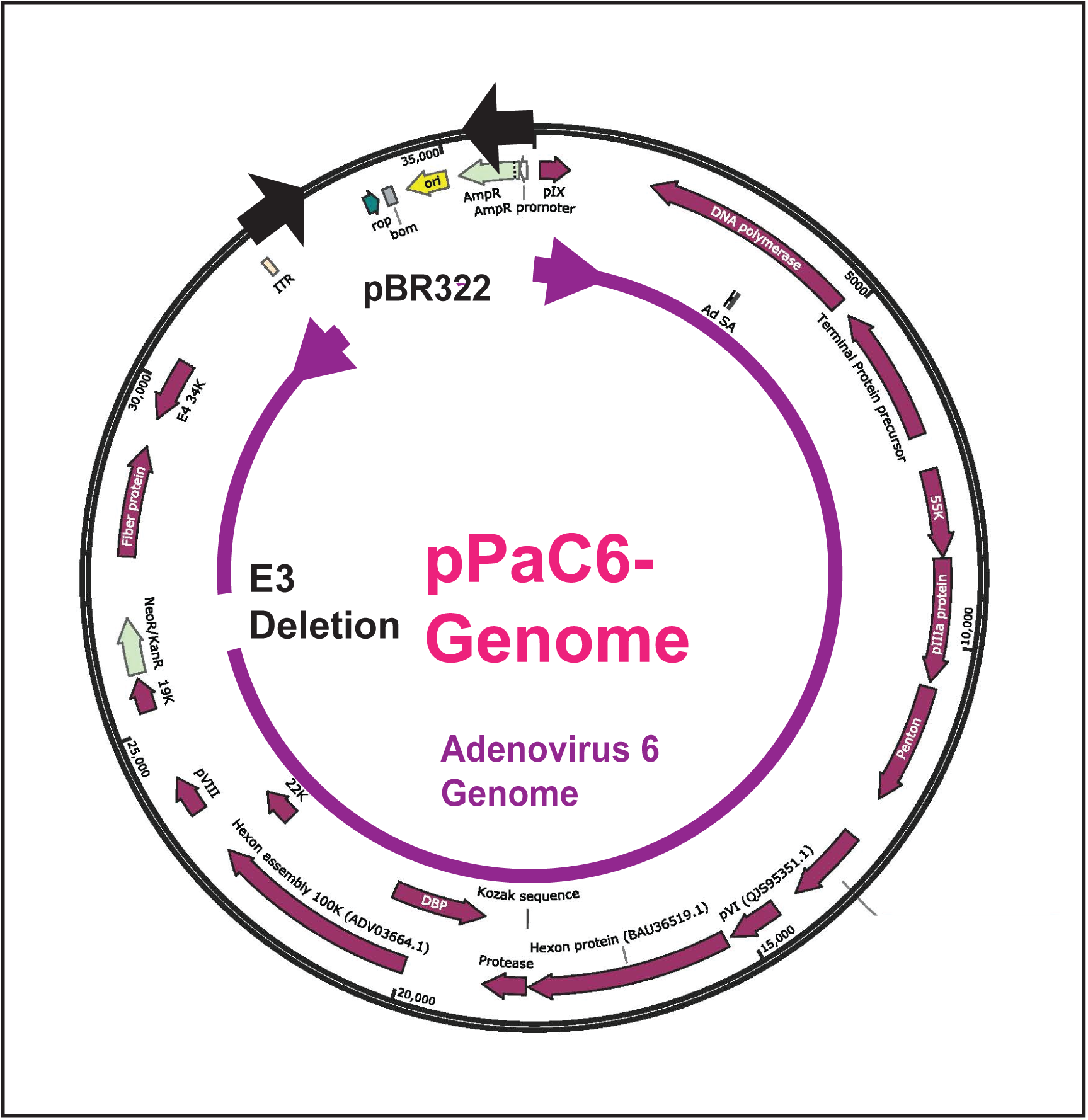
– The Genome of the pPaC6 Packaging Expression Plasmid. The pPaC6 plasmid carries a partial genome of an Adenovirus of the human serotype 6 in a pBR322-based plasmid. AmpR: ampicillin resistance gene; ITR: Adenovirus inverted terminal repeat; E3 deletion: partial deletion of the Adenovirus E3 region; E4 34K: Adenovirus gene E34K; NeoR/KanR: Neomycin/Kanamycin resistance gene; ori: origin of replication; pVI: Adenovirus gene pVI; pVIII: Adenovirus gene pVIII; pIX: Adenovirus gene pIX; rop: repressor of primer; 22K: Adenovirus gene 22K; 55K: Adenovirus gene 55k;

### Animal Challenge Studies

The present studies aimed to establish the efficacy of the GreFluVie6 vaccine in protecting animals against lethal challenges with an H5N1 pandemic avian influenza strain. Two animal species were chosen: (i) the mouse, for the broad availability of testing reagents, and (ii) the ferret, for its human-type infection of the respiratory tract (Belser et al., 2011). In contrast to some influenza strains, the wild-type H5N1 avian influenza strain *A/Vietnam/1203/2004* causes a disease in mice and ferrets without need of an adaption to the two animal species. Similarly to the human situation, clinical signs appear within 2 to 3 days after the viral challenge. Symptoms of lethargy, weight loss, ruffled fur and respiratory distress that would lead to death are prevalent in both species.

### Mouse Study

Based on our previous findings, an *i.m.* immunization route was chosen (Qi et al., 2024). The vaccine was delivered in a short-term prime/boost schedule (**Figure 2**). Mice were distributed into two groups that received the GreFluVie6 vaccine at doses of 1.5 x 10^7^ (**Medium**) and 5 x 10^6^ (**Low**) genome equivalents (GE). The **Negative Control** animals were injected with the Vector Formulation Buffer (VFB). Both cellular and humoral immune responses were evaluated. Four weeks after the second vaccine dose had been administered, mice were exposed to a lethal dose of the *A/Vietnam/1203/2004* virus given via an *i.n.* instillation. Four days thereafter, a subset of mice was euthanized. Their lungs were examined for the presence of live virus. For the remainder of the mice, body weights and health status were determined for twenty-one days.

**Figure 2.**
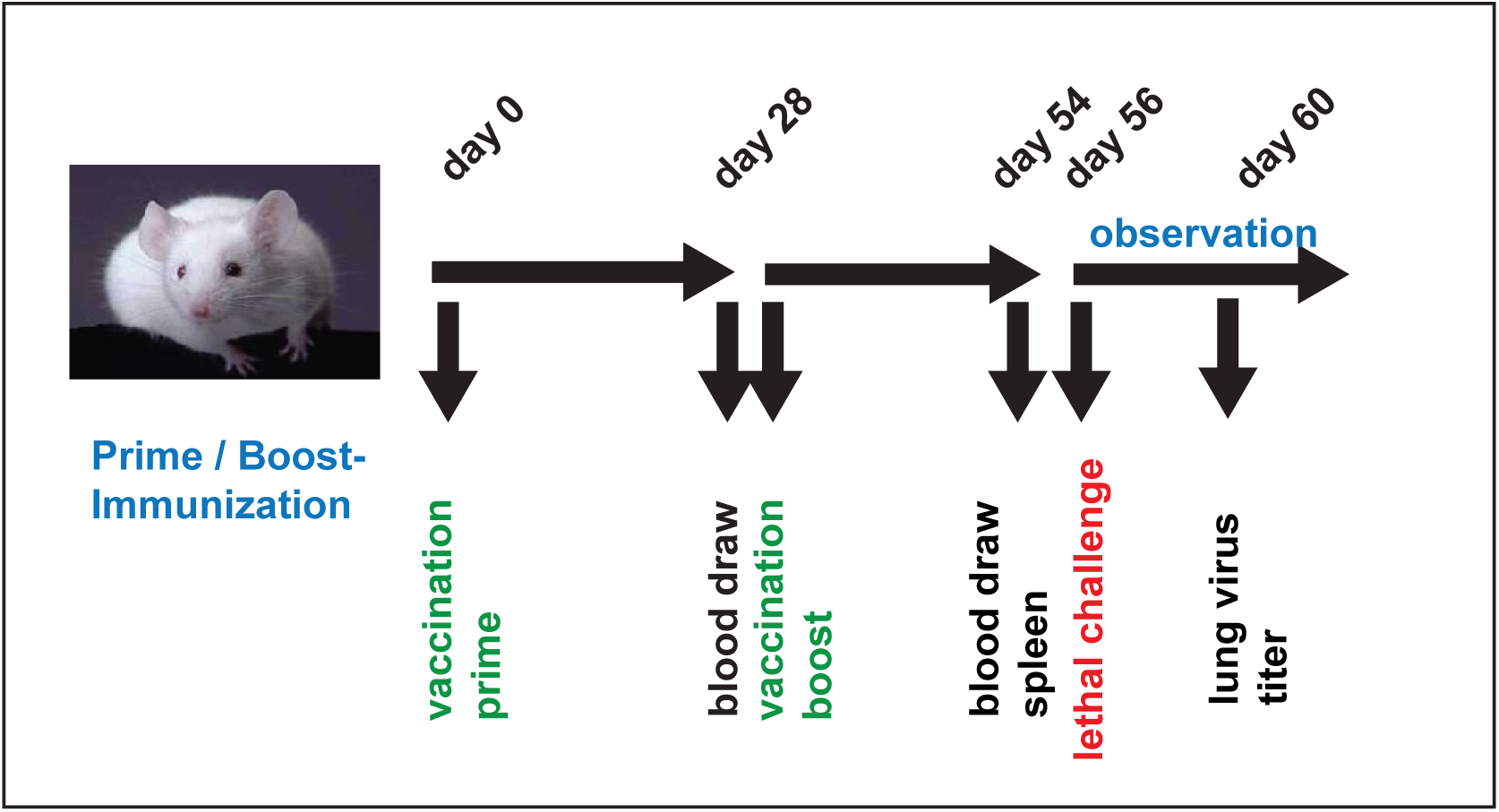
– Mouse Lethal Challenge Study. Balb/c mice were vaccinated by the *i.m.* route on day 1 and 27. They received a lethal challenge with the wild-type *A/Vietnam/1203/*2004 on day 29+/-1. They were observed through day 15-post challenge.

#### Health Status

**Figure 3** shows Kaplan-Meier survival curves and mean body weight changes for the different study groups. None of the **Negative Control** mice survived the lethal challenge (**Figure 3A**). They quickly lost body weight, and their health status deteriorated in parallel (**Figure 3A** and **B**). All **Medium** dose animals were completely protected with only minor weight losses of < 3%. They also did not show any negative health effects (**Figure 3C** and **D**). The **Low** dose still shielded 70% of the mice with an average body weight loss for the surviving animals of about 5% (**Figure 3E**). The affected animals quickly lost body weight, and their health deteriorated (**Figure 3E and F**).

**Figure 3.**
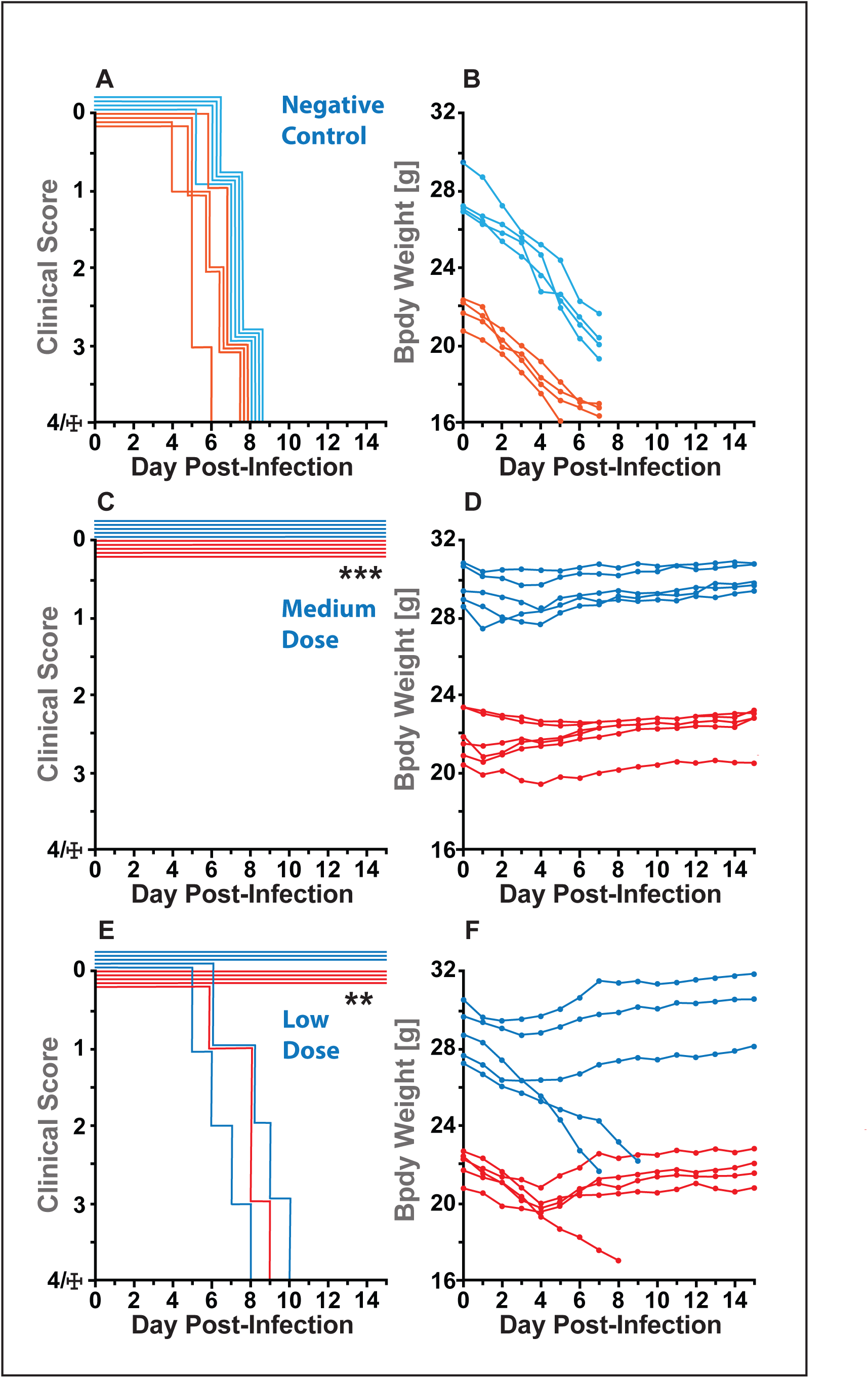
– Health Status, Survival and Body Weight of Mice After Lethal Challenge. Male (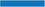) and female (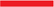) mice received the Vector Formulation Buffer (Negative Control) (A and B), or the GreFluVie6 vaccine at doses of 1.5 x 10^7^ GE (Medium dose) (C and D) and of 5 x 10^6^ GE (Low dose) (E and F) per *i.m.* injection. Their health and survival (A, C and E), as well as their body weights were surveilled (B, D and F. ** p<0.01;*** p<0.001

#### Virus Lung Titers

On day 4 post-challenge, two male and two female mice of each group were euthanized. Their lungs were removed and tested for the presence of infectious virus (**Figure 4**). **Negative Control** animals had high virus loads. Titers of virus in lungs of mice vaccinated with GreFluVie6 were significantly reduced in mice that received either **Medium** dose (p < 0.01) or **Low** dose (p < 0.05) vaccine.

**Figure 4.**
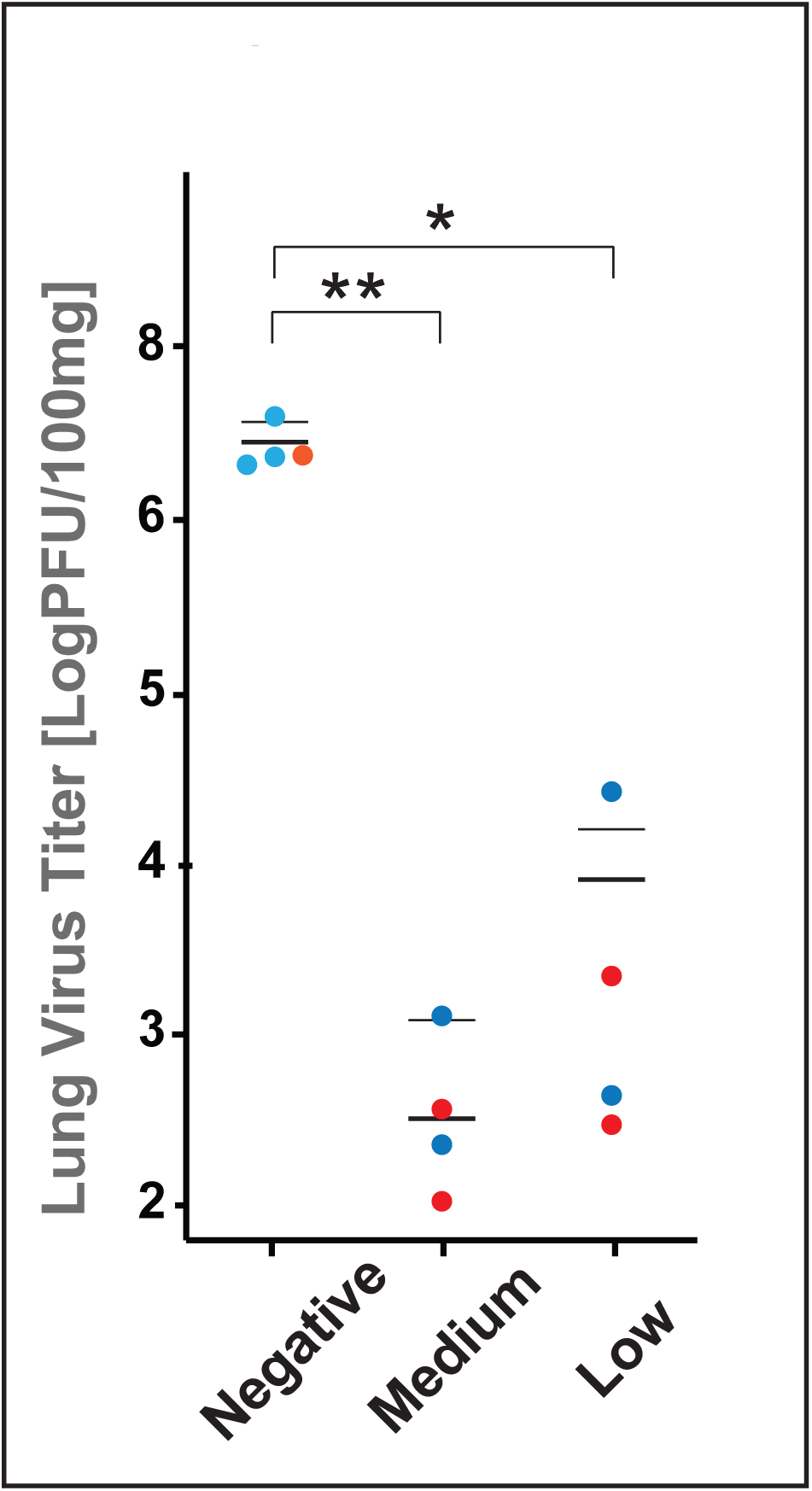
– Lung Virus Titers of Lethally Challenged Mice. On day 4 post challenge, two male (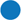) and two female (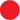) mice of each group that had received the Vector Formulation Buffer (Negative Control) or that had been immunized with the GreFluVie6 vaccine at doses of 1.5 x 10^7^ GE (Medium dose) and of 5 x 10^6^ GE (Low dose) per injection were sacrificed. Their lungs were harvested, and live virus titers were determined. The number of live viruses per 100 mg of lung tissue homogenate is depicted. * p<0.05; ** p<0.01;

#### Humoral Immune Responses

Blood was collected from mice 28 days after the second vaccine dose, but prior to the lethal challenge. Serum antibody titers were determined against the H5 hemagglutinin and N1 neuraminidase vaccine antigens that were encoded within the GreFluVie6 vaccine as well as another H5 variant. Specific antibodies were not detected in the sera of the **Negative Control** animals (**Figure 5A and B**). Mice that had been immunized with the **Medium** dose had raised antibodies to the vaccine H5 protein of *A/Vietnam/1203/2004* virus (**Figure 5A**). The seroconversion rate was 77.8%. When the same sera were tested against the H5 protein of the clade 2.1.3.2 *A/Indonesia/05/2005* H5 hemagglutinin, potent cross-reactivities were observed with a with a similar seroconversion rate (**Figure 5B**). Further analysis of the data revealed that all animals whose sera reacted with the vaccine H5 protein also had antibodies to the *A/Indonesia/05/2005* hemagglutinin (*data not shown*). Reducing the vaccine dose to **Low**, the seroconversion rate fell to 38.9% and 33.3% for the *A/Vietnam/1203/2004* and *A/IndonesiaI/05/2005* H5 hemagglutinins, respectively (**Figure 5A and B**). Little cross-reactivity was seen towards the H1 hemagglutinin of the *A/California/04/2009* at any vaccine dose (**Figure 5C**). Previously, we had determined that the GreFluVie6 vector drove the surface expression of the transgenes H5 and N1 by transduced cells at comparable levels (*data not shown*). However, when we tested the mouse sera for responses towards the N1 neuraminidase, we only detected low reactivities (**Figure 5D**). Here the seroconversion rate was reduced to 27.7%, and 22.2% for the **Medium** and **Low** vaccine doses, respectively. Yet, in ferrets, serum antibody responses to the N1 protein were strong (*see below*). The genome of GreFluVie6 vaccine is fully deleted of all endogenous Ad genes. It was packaged in an Ad capsid of the rare human serotype 6. We detected serum antibodies in 6 of the 12 mice in the **Medium** dose group (**Figure 5E**). However, the immune responses of this group did not reach significance when compared to the ones seen with the **Negative Control** animals. With the reduction in the vaccine dose, the immune responses vanished (**Figure 5E**). We also investigated neutralizing antibodies at the **Medium** vaccine dose with the help of a pseudotyped lentivirus. At this dose, the neutralizing antibody titers were comparable to the ones measured by ELISA (**Figure 5F**).

**Figure 5.**
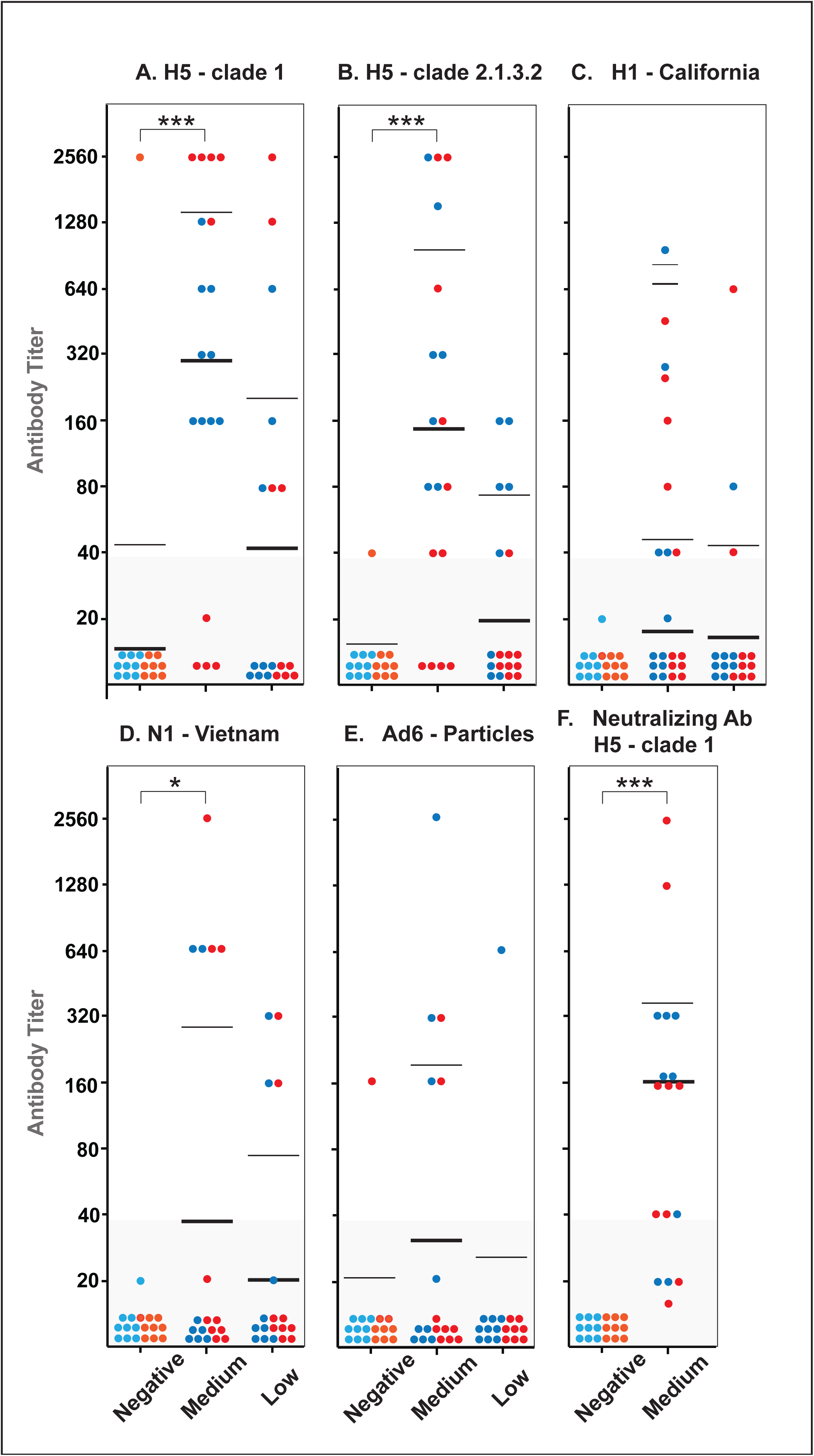
– Humoral Immune Responses of Vaccinated Mice. Male (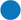) and female (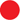) mice that had received the Vector Formulation Buffer (Negative Control) or that had been immunized with the GreFluVie6 vaccine at doses of 1.5 x 10^7^ GE (Medium dose) and of 5 x 10^6^ GE (Low dose) per *i.m.* injection were bled 28 days after the second immunization, but prior to the lethal viral challenge. Their sera were tested by ELISA for humoral immune responses to the H5 hemagglutinins of the *A/Vietnam/1203/*2004 virus (A) and *A/Indonesia/05/*2005 virus (B), the H1 hemagglutinin of the *A/California/04/*2009 virus (C), the N1 neuraminidase of the *A/Vietnam/1203/*2004 virus (D) and viral particles of the Adenovirus of the human serotype 6 (E). In addition, their ability of inhibiting cellular infections with a lentivirus pseudotyped with H5 hemagglutinin of the *A/Vietnam/1203/*2004 virus was examined (F). * p<0.05; *** p<0.001;

#### Cellular Immune Responses

After the second vaccination, but prior to challenge, the draining lymph nodes and spleens were harvested from two male and two female animals of each group. The T cells were tested for responses to the *A/Vietnam/1203/2004* H5 hemagglutinin using EliSpot assays. As summarized in **Figure 6A**, T cell immune responses to the H5 hemagglutinin protein were induced by the vaccine at both doses. They followed a TH1>TH2 response pattern characterized by a preferential release of interleukin-2 (IL-2), interferon-g (INF-g) and tumor necrosis factor-a (TNF-a) over interleukin-4 (IL-4), interleukin-6 (IL-6) and interleukin-10 (IL-10). To preferentially analyze the responses of the CD8^+^ T cells, we created a mixture of H5-derived peptides that carried an H-2K^d^-binding motif (Romero et al., 1991). When the T cells were challenged with these peptides, the responses followed similar TH1>TH2 responses (**Figure 6B**). The lower activity level may be due to the fact that the peptides used only bound to one of the three MHC class I molecules found in Balb/c mice (Mellor, 1986).

**Figure 6.**
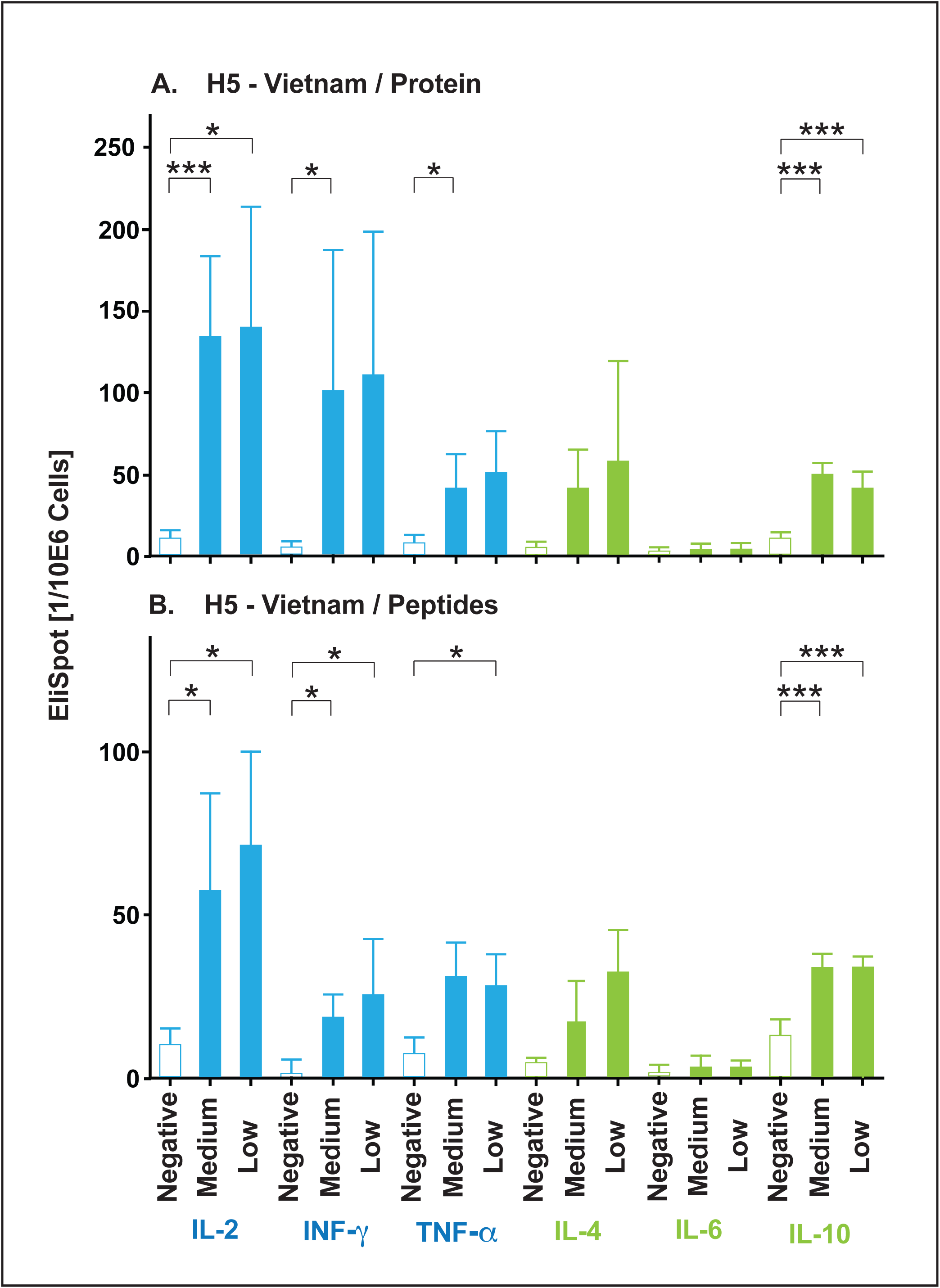
– Cellular Immune Responses of Vaccinated Mice. Two male and two female mice each group that had received the Vector Formulation Buffer (Negative Control) or that had been immunized with the GreFluVie6 vaccine at doses of 1.5 x 10^7^ GE (Medium dose) and of 5 x 10^6^ GE (Low dose) per *i.m.* injection were sacrificed 28 days after the second immunization, but prior to the lethal viral challenge. Their spleens and draining lymph nodes were harvested and stimulated *in vitro* with the H5 hemagglutinin of the *A/Vietnam/1203/*2004 virus (A) or a mixture of H5 hemagglutinin-derived peptides designed to bind to the H-2K^d^ MHC class I molecule (B). The number of EliSpots per 10^6^ lymphocytes detecting the release of IL-2, IFN-γ, TNF-α, IL-4, IL-6 and IL-10 is depicted. * p<0.05; *** p<0.001;

### Ferrets

The ferret studies followed a similar protocol. The three experimental groups consisted of the **Negative Controls** and animals that received the GreFluVie6 vaccine at **High** (3 x 10^9^ GE) or **Medium** (1 x 10^9^ GE) doses. The animals were immunized via the *i.m.* route with a prime/boost protocol. Four weeks after the last vaccination, the ferrets were challenged (> 200-fold LD50) with *A/Vietnam/1203/2004* virus given via an *i.n.* instillation. The animals were weighed initially daily and then every second day. On days 1, 3 and 5 after the virus inoculation, nasal flushes were performed, and titers of live virus were determined by plaque assay. Four days post challenge, two ferrets of each group were sacrificed to evaluate lung virus titers.

#### Health Status

The data for the ferrets’ health status, body weights and body temperatures are summarized in **Figure 7**. The health of the **Negative Control** animals deteriorated quickly (**Figure 7A**). The ferrets experienced significant body weight losses and suffered under elevated body temperatures (**Figure 7B and C**). They became moribund by day 8 and were euthanized. Both the **High** and **Medium** dose animals were completely protected against the lethal viral challenge (**Figure 7B and G**). Some animals showed slightly reduced activities after the exposure to the challenge virus. Significant changes in body weights were not observed (**Figure 7E** and **H**). The body temperatures of the vaccinated ferret increased post challenge, but normalized by day eight (**Figure F** and **I**).

**Figure 7.**
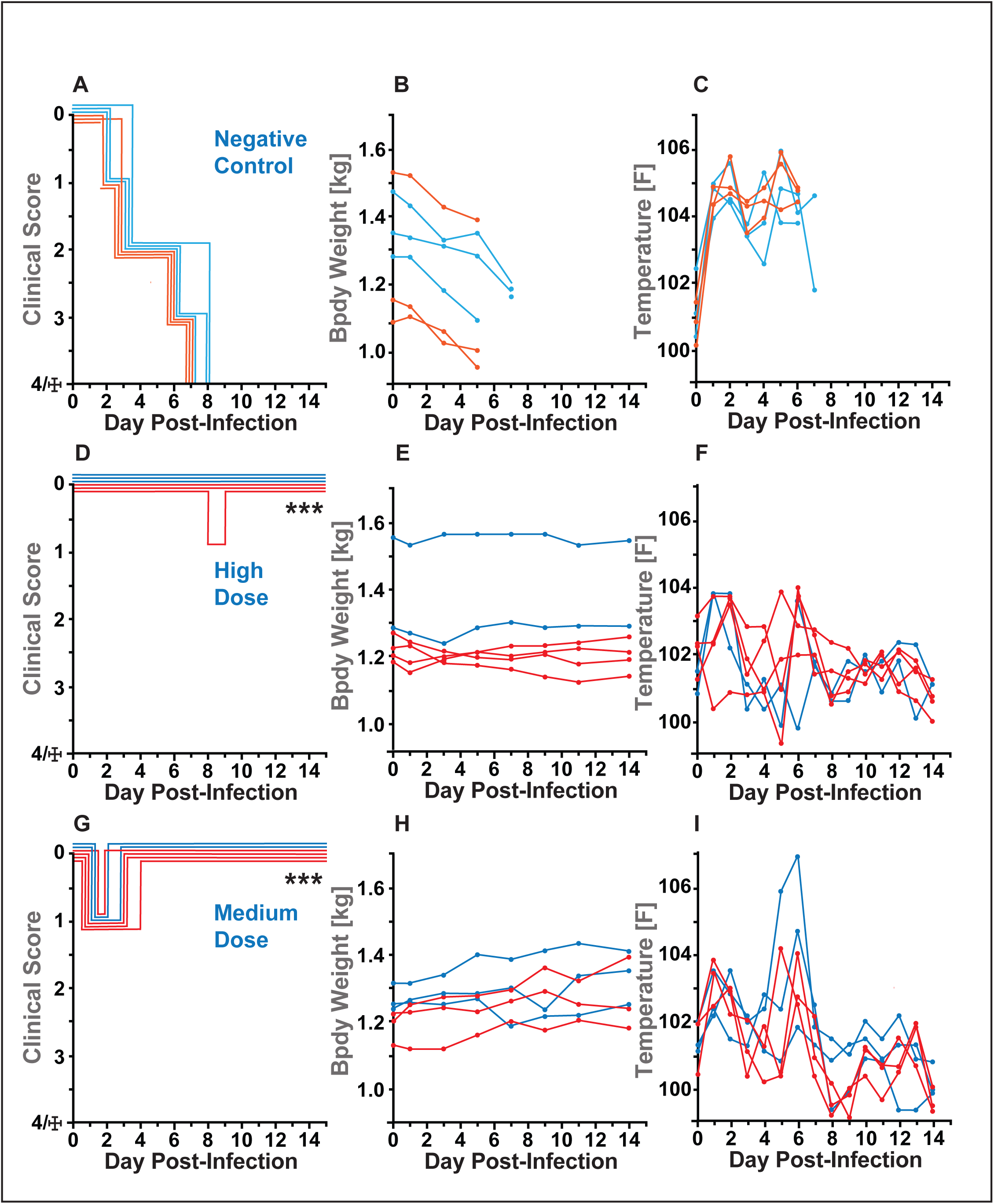
– Health Status, Survival and Body Weight of Ferrets After Lethal Challenge. Three male (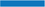) and three female (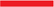) ferrets received the Vector Formulation Buffer (Negative Control) (A, B and C), or the GreFluVie6 vaccine at doses of 3 x 10^9^ GE (High dose) (D, E and F) and of 1 x 10^9^ GE (Low dose) (G, H and I) per *i.m.* injection. Their health and survival (A, D and G), as well as their body weights (D, E and H) and body temperatures (C, F and I) were surveilled for 15 days post challenge *** p<0.001

#### Virus Titers

Nasal flushing samples were collected on days 1, 3 and 5 post challenge. On day 1, the virus titers did not differ between the three experimental groups (**Figure 8A**). On day 3, the virus load increased for the **Negative Control** animals, whereas it started to decrease for the **High** and **Medium** vaccine dose ones (**Figure 8B**). On day 5, the virus levels showed a small reduction for the **Negative Control** animals. However, they were significantly lowered for all ferrets that had been vaccinated (**Figure 8C**). The **High** dose animals saw the most efficient reduction. Two animals of each group were euthanized on day 4 post challenge. Their lungs were harvested and homogenized. Live viruses were detected in both **Negative Control** animals. None was seen in any of the vaccinated ferrets (**Figure 9**).

**Figure 8.**
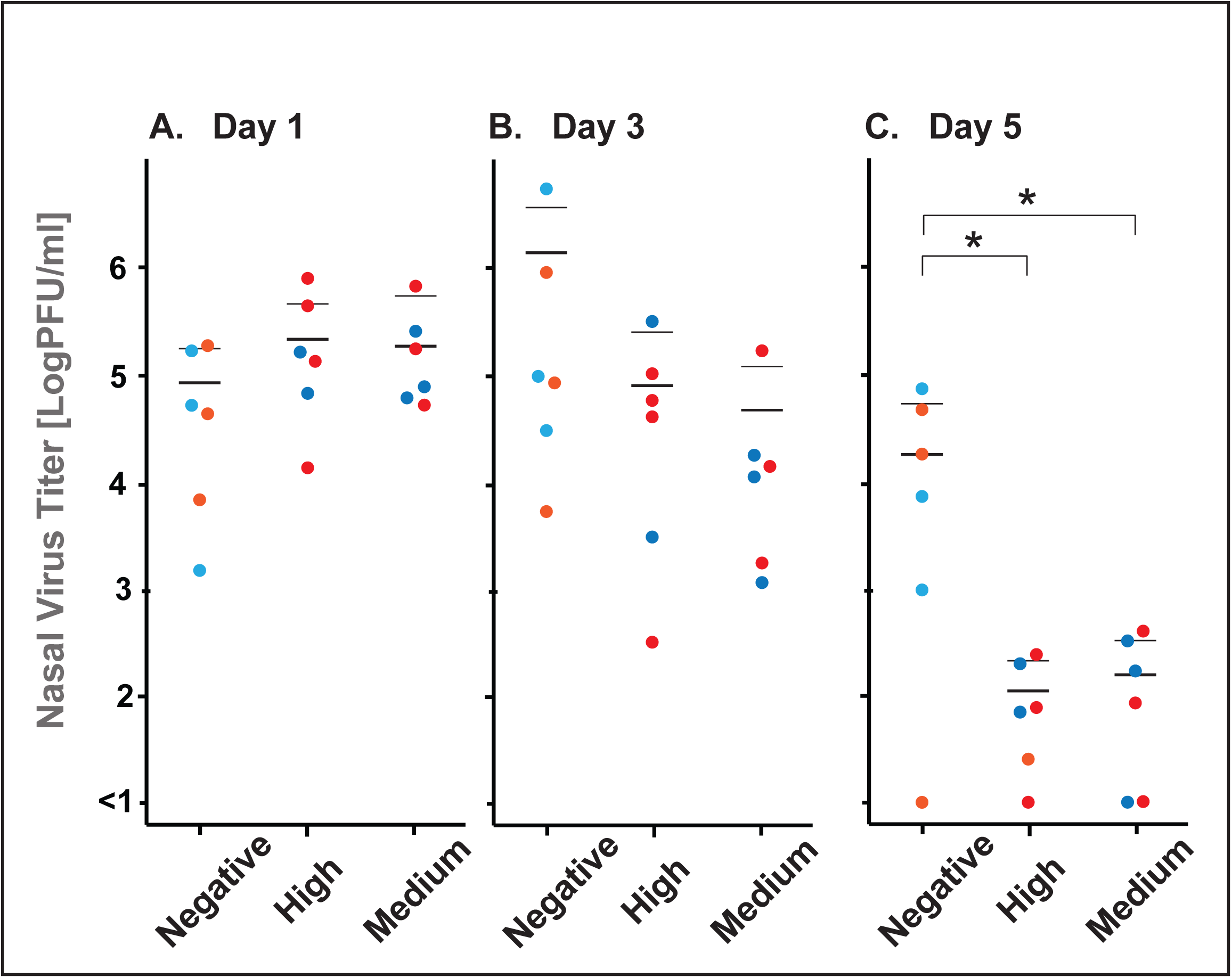
– Nasal Lavage Virus Titers of Lethally Challenged Mice. On days 1, 3 and 5 post challenge, nasal lavage samples of male (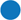) and female (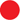) ferrets of each group that had received the Vector Formulation Buffer (Negative Control) or that had been immunized with the GreFluVie6 vaccine at doses of 3 x 10^9^ GE (High dose) and of 1 x 10^9^ GE (Low dose) per *i.m.* injection were collected. Live virus titers of the lavage sample were determined. The number of live viruses per 1 ml of material is depicted. * p<0.05;

**Figure 9.**
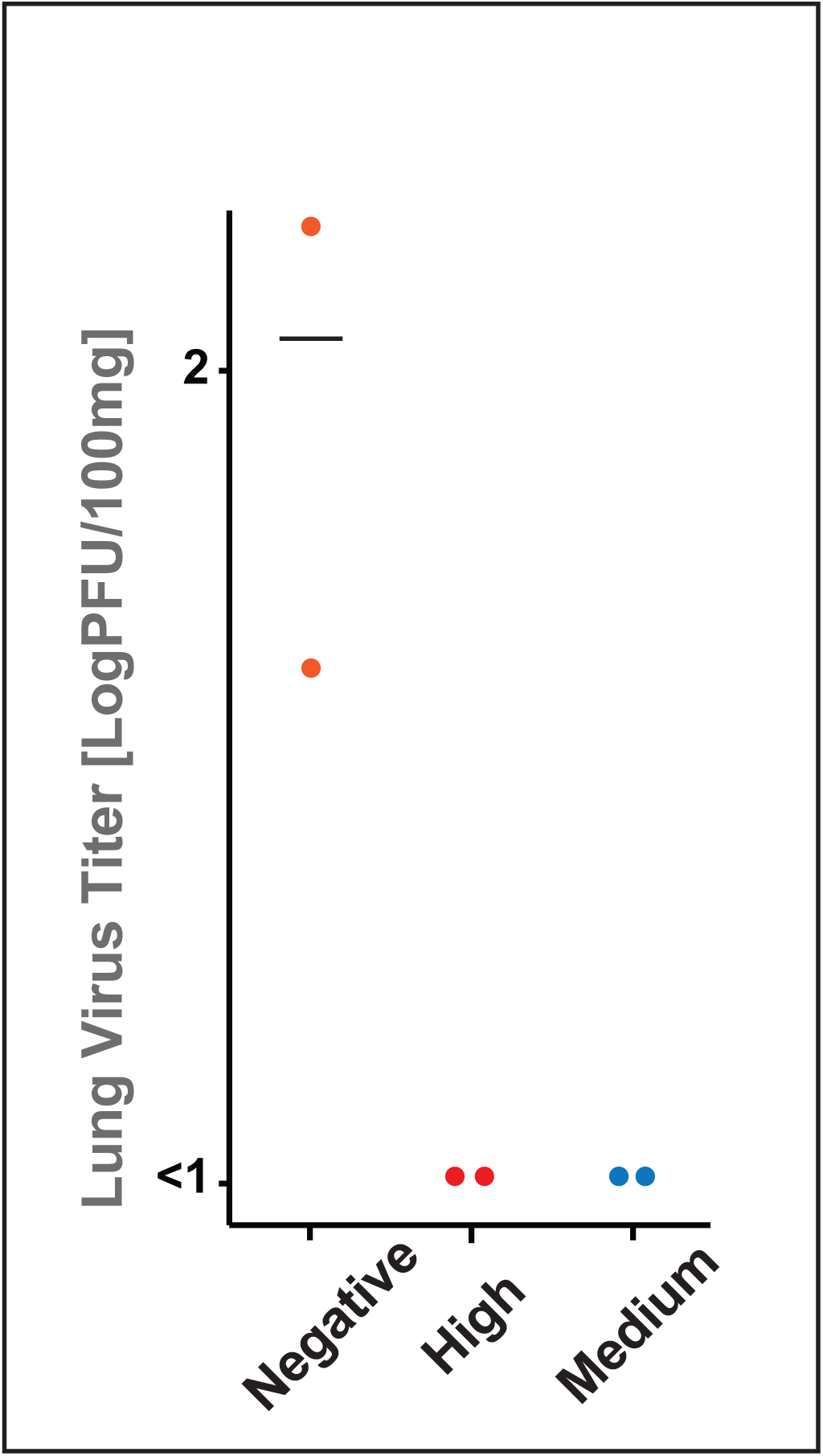
– Lung Virus Titers of Lethally Challenged Ferrets. On day 4 post challenge, one male (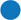) and one female (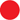) ferrets of each group that had received the Vector Formulation Buffer (Negative Control) or that had been immunized with the GreFluVie6 vaccine at doses of 1.5 x 10^7^ GE (Medium dose) and of 5 x 10^6^ GE (Low dose) per injection were sacrificed. Their lungs were harvested, and live virus titers were determined. The number of live viruses per 100 mg of lung tissue homogenate is depicted.

#### Humoral Immune Responses

Prior to the first and second vaccination and viral challenges, the animals were bled. Their sera were tested for binding to the H5 hemagglutinins and the N1 neuraminidase proteins encoded in the GreFluVie6 vaccine. Specific antibodies were not detected in the sera of any of the **Negative Controls** (*data not shown*) or of animals prior to vaccination (**Figure 10**). At the **High** dose, strong specific antibody responses were already induced to the vaccine H5 hemagglutinin of the *A/Vietnam/1203/2004* virus after the first immunization (**Figure 10A**). They were further enhanced by the second injection. In parallel, the seroconversion rate increased from 83% to 100%. At the **Medium** dose, ferrets only showed significant antibody titers after the booster injection, at which point all animals had seroconverted (**Figure 10A**). Similar response patterns were seen, when the sera were tested against the H5 hemagglutinin of the clade 2.1.3.2 *A/Indonesia/05/2005* virus (**Figure 10B**). All ferrets that had received the vaccine at the **High** dose responded strongly after the first dose and even more pronounced after the second one. Reducing the vaccine dose to **Medium** resulted in potent responses only after the vaccine boost. When we tested the animal sera against the H5 of a clade 2.3.4.4b influenza virus (*A/Chicken/Netherlands/14015526/2014*), we observed potent cross-reactivities. The serum antibodies of both the **High** and **Medium** dose groups showed some binding to this H5 protein after the prime dose (**Figure 10C**). The titers were further boosted with the second vaccination. Similarly to the mouse models, specific antibodies reactive with the H1 hemagglutinin (*A/California/04/2009*) were not detected (**Figure 10D**). In contrast to mice, ferrets raised significant humoral immune responses against the vaccine N1 neuraminidase protein (**Figure 10E**). At the **High** dose, specific antibodies became detectable after the first immunization. Their titers were further increased after the vaccine boost. **Medium** dose animals only raised significant responses after the second injection. The seroconversion rate ultimately reached 100% for both groups. When we examined the sera for the presence of antibodies specific for the Ad capsid, we only saw low responses in the **High** dose group and only after the second vaccination (**Figure 10F**). Vaccination with the **Medium** dose failed to raise significant antibody response to the vector. We studied virus neutralization using pseudotyped lentiviruses (Crawford et al., 2020; Ferrara & Temperton, 2018; Temperton et al., 2007). At the **High** dose, significant titers of antibodies able to neutralize *A/Vietnam/1203/2004* and *A/Indonesia/05/2005* H5 hemagglutinin-expressing particles were detected (**Figure 10G** and **F**). They were further boosted by a second vaccine dose resulting in 100% seroconversion. Animals that had received the vaccine at the **Medium** dose only raised significant neutralizing antibody responses after the second dose (**Figure 10G**). They remained borderline when the *A/Indonesia/05/2005* H5 hemagglutinin was presented as target on the pseudovirus (**Figure 10H**).

**Figure 10.**
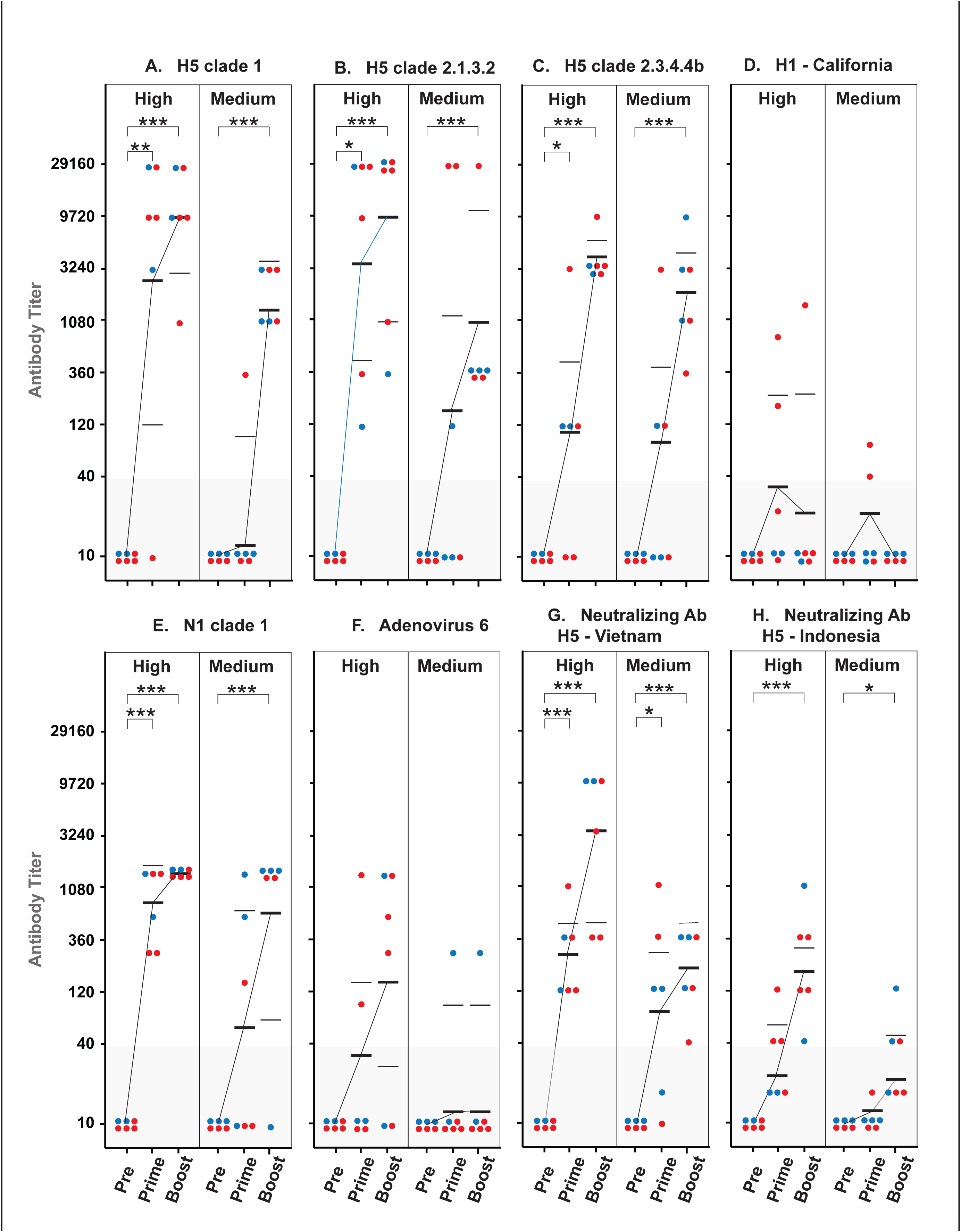
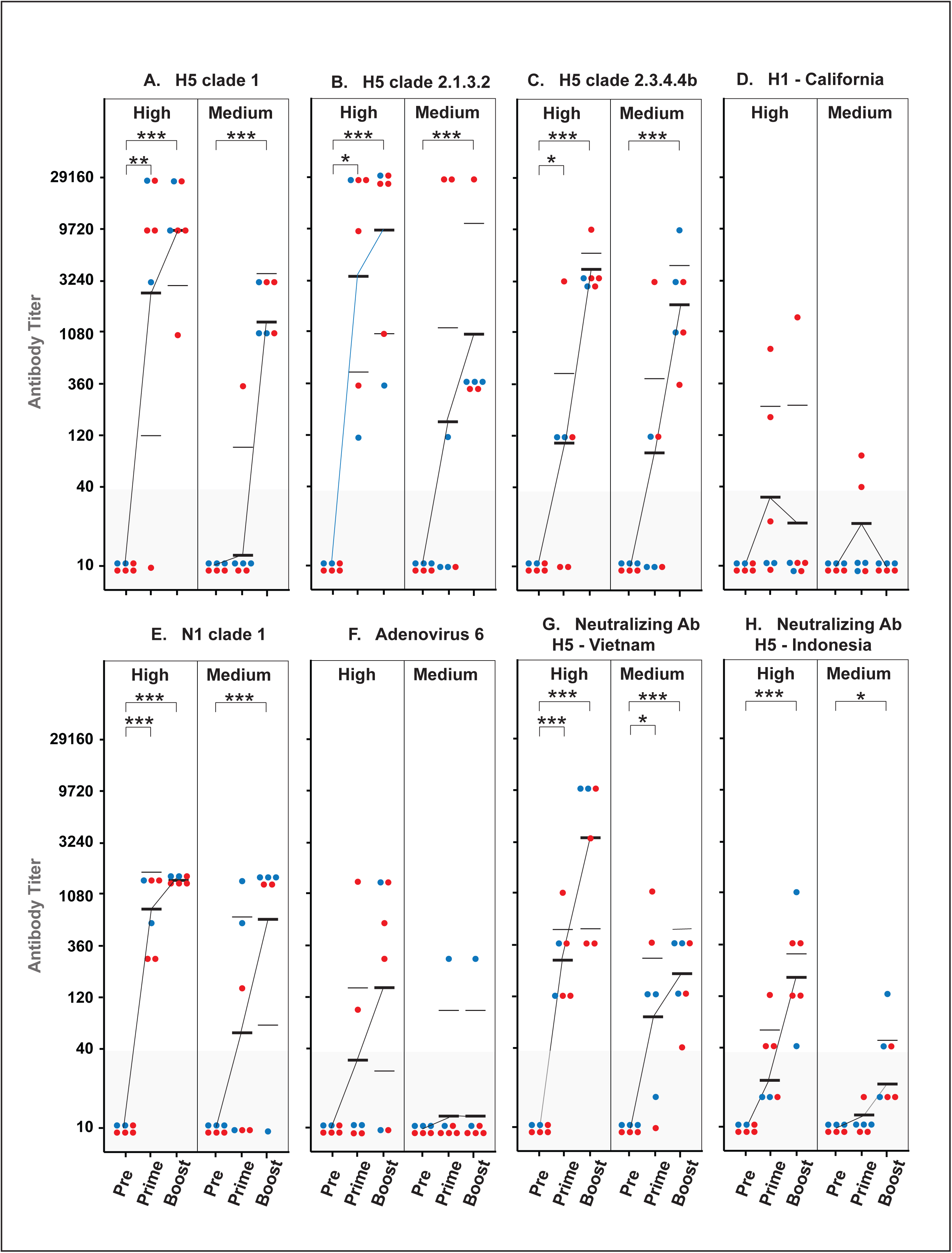
-Humoral Immune Responses of Vaccinated Ferrets. Male (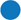) and female (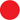) ferrets that had received the Vector Formulation Buffer (Negative Control) or that had been immunized with the GreFluVie6 vaccine at doses of 3 x 10^9^ GE (High dose) and of 1 x 10^9^ GE (Medium dose) per *i.m.* injection were bled 28 days after the second immunization, but prior to the lethal viral challenge. Their sera were tested by ELISA for humoral immune responses to the H5 hemagglutinins of the *A/Vietnam/1203/*2004 virus (A), *A/Indonesia/05/*2005 virus (B) and *clade 2.3.4.4b* (C), the H1 hemagglutinin of the *A/California/04/*2009 virus (D), the N1 neuraminidase of the *A/Vietnam/1203/*2004 virus (E) and viral particles of the Adenovirus of the human serotype 6 (F). In addition, their abilities of inhibiting cellular infections with a lentivirus pseudotyped with H5 hemagglutinin of the *A/Vietnam/1203/*2004 (G) and *A/Indonesia/05/*2005 (H) viruses were examined (F). * p<0.05; ** p<0.01;*** p<0.001;

## DISCUSSION

The present work builds on the results of an earlier proof-of-principle study, in which we had demonstrated that the GreVac platform of a 4^th^ generation fully deleted helper virus-independent Ad vector could be used to create a potent vaccine (Qi et al., 2024; Qi et al., 2024). In these earlier experiments, the same Ad genome that carried a transgene expression cassette for the H5 hemagglutinin and the N1 neuraminidase had been packaged into an Ad capsid of the human serotype 5 (Ad5) (Qi et al., 2024). We showed that a GreVac-based vaccine fully protected mice against a lethal challenge with the wild-type *A/Vietnam/1203/2004* virus and that *i.m.* injections proved more efficient than *s.c.* and *i.n.* vaccination routes contrary to findings published by others (Casimiro et al., 2003; Chen et al., 2024; Coughlan et al., 2018; Happe et al., 2024; Jia et al., 2024; Jin et al., 2022; Li et al., 2023; Tapia et al., 2016).

Infections with Ad of the human serotype 5 are frequent occurrences in the human population (Akello et al., 2020). Therefore, pre-existing immunities are quite common that interfere with activities of Ad5-based vectors both of early-generation and fully deleted design (Fausther-Bovendo & Kobinger, 2014). Switching to an Ad vector system of the rare human serotype 6 (Ad6) will in most cases (> 95%) prevent an interference by an established immune response (Akello et al., 2020). Another issue of early-generation Ad vectors is caused by their delivery of numerous endogenous Ad genes in concert with the vaccine antigens (Sayedahmed et al, 2022). Potent anti-Ad responses are raised that in turn can affect the efficacy of a second application of the same vector (Ahi et al., 2011). The GreFluVie6 vector is based on the fully deleted helper virus-independent GreVac vaccine platform. The immune target size and therefore its immunogenicity are reduced as seen in **Figures 5E** and **10F**. Consequently, efficient short-term prime/boost vaccination were possible both in the mouse and the ferret with the same vaccine.

We were able to directly study protection against viral infections with the wild-type *A/Vietnam/1203/2004*. Adaptation of this influenza virus was not required for the mice or ferrets. In both models, we identified vaccine doses that delivered complete protection against lethal challenges with the *A/Vietnam/1203/2004* virus (**Figure 3** and **7**). In the mouse, they corresponded to approximately 2% of ones previously used with early generation Ad vectors (Gao et al., 2006; Kobinger et al., 2007; Patel et al., 2010; Roy et al., 2007). Decreasing the vaccine dose in the mouse from **Medium** to **Low** reduced the protection rate from 100% to the still highly positive 70%. In the ferret, full protection was achieved with both the **High** and **Medium** vaccine doses. In both models, significant body weight losses and deterioration of the animals’ health were prevented. In the ferret, initial increases in the body temperatures normalized by day 8 post challenge. Elevated temperatures were more pronounced for animals of the **Medium** dose group. This fact might indicate that the immune system required an extended ramp-up. This delay of response might have been revealed by the very high challenge virus dose of > 200-fold LD50.

Both in the mouse and the ferret, *i.m.* vaccinations with GreFluVie6 minimized the presence of live virus in the lung in a dose dependent way (**Figure 4** and **9**). In the ferret, both vaccine doses significantly reduced virus titers in the nasal cavity. It could therefore be concluded that the vaccination with GreFluVie6 would not only protect the individual, but also limit viral transmissions within a population.

The GreFluVie6 induced strong immune responses. In the mouse, prime/boost immunization produced high levels of serum antibodies that bound to the clade 1 *A/Vietnam/1203/2004* H5 hemagglutinin (**Figure 5**). Potent cross-reactivities towards the clade 2.1.3.2 *A/Indonesia/05/2005* H5 hemagglutinin were observed. Similar response patterns were seen for the ferrets (**Figure 10**). A strong reactivity to the vaccine clade 1 H5 hemagglutinin was mated to a potent cross-reactivity to the clade 2.1.3.2 H5 hemagglutinin. Expanding these studies further to the clade 2.3.4.4b-type H5 hemagglutinin revealed even broader cross-reactivities of the vaccine induced humoral responses. Using H5 pseudotyped lentiviruses, neutralizing antibody titers were comparable to the ones measured by ELISA (**Figure 5** and **10**). When the higher vaccine doses were used, we observed significant activities in both animal models already after the first vaccination. When the vaccine was dosed at the lower levels, strong neutralization required a boost. No serum antibodies were detected that bound to the H1 hemagglutinin in either animal model.

In test cells transduced with the GreFluVie6 vector, the H5 and N1 proteins were expressed at similar levels (*data not shown*). However, the mice failed to raise significant levels of reactive antibodies (**Figure 5**). In contrast, ferrets raised strong immune responses to the N1 antigen (**Figure 10**). This finding confirmed that the N1 protein in the GreFluVie6 genome was indeed expressed in animals. The ability of Balb/c to raise potent humoral immune responses to N1 neuraminidase might have been limited by the low vaccine doses used in these studies.

The mouse model offered the opportunity to investigate the activity of T cells. In contrast to humoral immune responses, the cellular ones seemed less dose dependent (**Figure 6**). When tested against the *A/Vietnam/1203/2004* H5 hemagglutinin antigens, T cell activities were comparable at both vaccine doses. They were both biased towards TH1-type and against TH2-type responses. To better evaluate the specific activity of CD8^+^ T cells, we challenged them with a mixture of H5-derived peptides that carried H-2K^d^-binding epitopes. The overall response pattern, *i.e.* TH1>TH2, remained similar albeit at lower levels. These findings suggested that T cells were more sensitive to the vaccine, yet potent humoral immune responses were required to deliver full protection.

GreVac-based GreFluVie6 vaccine is completely deleted of all Ad genes. It was engineered as a 4^th^ generation of Ad vectors that was produced in the absence of and without the contaminations with helper viruses (Dormond et al., 2009, 2010). The only Ad-derived antigens are delivered as proteins of the Ad vector particle. The absence of endogenous Ad genes severely limited the induction of anti-Ad vector immune responses in both mice and ferrets (**Figure 5** and **10**). We were therefore able to employ short-term prime-boost vaccination protocols in contrast to some early generation Ad vectored vaccine that had resorted to vaccines of a completely different design for the second dose (Casimiro et al., 2003; Chen et al., 2024; Coughlan et al., 2018; Happe et al., 2024; Jia et al., 2024; Li et al., 2023; Tapia et al., 2016). The ability of GreVac vaccines to minimize induction of interfering immune responses might explain why we were able to succeed with significantly lower than the ones that had been reported with vaccine based on early generation Ad vectors (Gao et al., 2006; Morse et al., 2013; Patel et al., 2010; Singh et al., 2008). This increased potency might also explain the cross-reactive immune responses towards H5 hemagglutinins of both clade 1, clade 2.1.3.2 and clade 2.3.4.4b avian influenzas that were elicited by the GreFluVie6 vaccine.

## MATERIAL AND METHODS

### Animals

Male and female 6 to 8 weeks-old *BALB/c* mice were obtained from Charles River Laboratories (Wilmington, MA). The mice were quarantined for 7 days before use. Male and female 12 week-old influenza-free *Mustela putorius furo* ferrets were obtained from Triple F Farms (Gilette, PA). They had been castrated or spayed. The ferrets were quarantined for 7 days before use. All animals were maintained at BSL2 (vaccination phase) or BSL3 (challenge phase) facilities at Colorado State University

### Challenge Virus

The influenza virus *A/Vietnam/1203/2004* (H5N1) was obtained from the Centers for Disease Control (Atlanta, GA). Viral propagation and titration were performed as previously described (Qi, 2024).

### Vaccine

GreFluVie6 (H5N1) vaccine was produced and purified as described in (Qi et al., 2024). It was diluted at concentrations of 3 x 10^8^ GE/ml (**Medium** dose) or 1 x 10^8^ GE/ml (**Low** dose) for the mouse studies, and at concentrations of 6 x 10^9^ GE/ml (**High** dose) or 2 x 10^9^ GE/ml (**Medium**) for the ferret studies. The vaccine was suspended in vector formulation buffer (VFB, Tris HCl 20mM pH7.8, NaCl 140mM, MgCl2 5mM, EDTA 0.5mM, polyethylene glycol 4000 (PEG 4000) 0.02 mM, and sucrose 2% (W/V)). Samples of the VFB were used as **Negative Control**.

### Animal Studies

Groups of mice (9 males and 9 females per group) were vaccinated by the *i.m.* route in 50 μl of volume on two occasions, day 1 and 27, with each vaccine dose containing 1.5 x 10^7^ GE (**Medium** dose) of 5 x 10^6^ GE (**Low** dose) of the GreFluVie6 vaccine per injection. Groups of 8 ferrets (4 males and 4 females per group) received 3 x 10^9^ GE (**High** dose) or 1 x 10^9^ GE (**Medium** dose) of the GreFluVie vaccine per 500 μl *i.m.* injection. **Negative Control** mice and ferrets received 50 μl or 500 ml of VFB by *i.m.* injections, respectively. For influenza virus challenges, mice were anesthetized by intraperitoneal injection of ketamine/xylazine (50 mg/kg/5 mg/kg) prior to the *i.n.* challenge with a 90-μl suspension of the wild-type influenza *A/Vietnam/1203/2004* virus corresponding to 20-fold LD50. Ferrets received the *i.n.* challenge with a 200-ml suspension of the wild-type influenza *A/Vietnam/1203/2004* virus corresponding to 200-fold LD50. All animals received the *A/Vietnam/1203/2004* challenge virus 29+/-1 days after the second vaccine dose. They were weighed prior to virus challenge and then daily or every other day thereafter. They were observed for morbidity and mortality through day 15 post-challenge. The body temperatures of ferrets were measured every second day.

### Live Virus Titer Determination

Lung homogenates or nasal lavage samples were centrifuged. Varying dilutions of the clarified supernatants were assayed in triplicate for infectious virus in MDCK cells, with virus titers calculated (Barnard et al., n.d.). Virus titers less than 200 were considered negative. Nasal flushes were obtained from ferrets under light ketamine-xylazine anesthesia. Briefly, 1 ml of cell culture medium with 1% BSA was dripped into the ferret’s nares, stimulating them to sneeze nasal secretions into a petri dish held next to the ferret’s head.

### Hemagglutinin, Neuraminidase and Adenoviral Particle-Binding Antibody ELISA

#### (A/Chicken/Netherlands/14015526/2014)

On day 1, 96-well plates (Corning, NY) were coated with anti-His-tag antibodies and stored overnight at room temperature (ThermoFisher, MA). The plates were washed and were blocked by a 5% BSA blocking buffer before the His-tagged proteins were added (H1 (*A/California/07/2009*) – Creative Biomart, NY: H5 (*A/Indonesia/5/2005*), H5 (*A/Vietnam/1203/2004*) - MyBiosource, CA; H5 clade 2.3.4.4b (*A/Chicken/Netherlands/14015526/2014*) – Native Antigen, Kidlington, UK). After an incubation at room-temperature, the free antigens were washed away, and the mouse or ferret sera were titrated into the wells. After the plates had been washed to remove the unbound antibodies, the extent of mouse antibody binding was developed with horseradish peroxidase-labeled goat anti-mouse Ig or anti-ferret Ig polyclonal antibodies (Southern Biotech, AL). The combinations of anti-H5 rabbit polyclonal antibodies and an HRP-tagged goat anti-rabbit Ig polyclonal antibody were used as positive control (Southern Biotech, AL). To evaluate antibody responses against Adenovirus 6 proteins, fixed Adenovirus virus paticles were captured onto the well surface with a goat anti-Adenovirus polyclonal antibody (MyBiosource, CA). The N1 antigen (R&D Systems, MN) was used to directly coat the wells, before the mouse or ferret sera were added.

### ELISPOT Assays of Cellular Immune Reponses of Vaccinated Mice

Draining lymph nodes and spleens were harvested from 2 female and 2 male mice of each group prior to the lethal viral challenge. The number of harvested cells was determined. 1 x 10^5^ cells were added to wells of 96-well plates of EliSpot. They were stimulated *in vitro* with the H5 hemagglutinin of the *A/Vietnam/1203/2004* virus (……) or a mixture of H5-derived peptides with binding motives to the H-2K^d^ MHC class I molecule (LYQBPTTYI; LYDKVRLQL; IYSTVASSL) (Romero et al., 1991). Following the manufacturers protocols, the cultures were developed for the release of interleukin-2 (IL-2), interleukin-4 (IL4), interleukin-6 (IL-6), interleukin-10 (IL-10), g-interferon, and tumor necrosis factor-a.

### Virus Neutralizing Antibody Assay

The hemagglutinin genes derived from *A/Vietnam/1203/2004* and *A/Indonesia/5/2005* were moved into the multiple cloning site of the pLVX-IRES-ZsGreen1 plasmid behind a CMV promoter (Takara Bio, San Jose, CA). The pseudotyped lentivirus was produced using the Lenti-X HTX packaging system (Takara Bio, San Jose, CA). The virus titers were determined using the Lenti-X CoStix Plus method (Takara Bio, San Jose, CA). The virus multiplicity of infection was determined that resulted in the transduction > 95% Q7 cells in test cultures. The Q7 cells were seeded on 96-well plates at 1 x 10^4^ cells per well in IMDM containing 5% FBS (Hyclone) 24 hours prior to use. On the next day, serial 2-fold dilutions of serum samples from five mice were prepared in serum-free medium starting at 1:5 dilution and ending at 1:4860. Each serum dilution was mixed 1:1 (0.1 ml) with serum-free media containing pseudotyped lentiviruses. After incubation at room temperature for 1 h, the serum-influenza virus mixture (0.2 ml) was transferred to a well containing the Q7 cells and incubated for 2 days. The cells were released from the well using a mixture of Trypsin and EFTA. The extent of their fluorescence of 50 cells per sample was analyzed under a fluorescence microscope using imageJ.net software. A neutralizing antibody titer was assigned when the overall fluorescence intensity had reached a level of ≥ 50%.

### Statistical Analysis

Kaplan-Meier survival curves were generated and compared by the Log-rank (Mantel-Cox) test followed by pairwise comparison using the Gehan-Breslow-Wilcoxon test in Prism 6.0f (GraphPad Software Inc., La Jolla, CA). The mean body weights were analyzed by analysis of variance (ANOVA) followed by Tukey’s multiple comparison test using Prism 6.0f. Virus titer differences were evaluated by ANOVA on log-transformed values assuming equal variance and normal distribution. Following ANOVA, individual treatment values were compared to placebo control by Tukey’s pair-wise comparison test using Prism 6.0f. The results from virus neutralization assays and hemagglutination inhibition assays were analyzed by ANOVA followed by Tukey’s multiple comparison test using Prism 6.0f.

### Ethics Regulation of Laboratory Animals

This study was conducted in accordance with the approval of the Institutional Animal Care and Use Committee of Colorado State University. The work was done in AAALAC-accredited Colorado State University. The U. S. Government (National Institutes of Health) approval was maintained in accordance to the latest National Institutes of Health Guide for the Care and Use of Laboratory Animals.

## REFERENCES

Abbink, P., Larocca, R. A., De La Barrera, R. A., Bricault, C. A., Moseley, E. T., Boyd, M., Kirilova, M., Li, Z., Ng’ang’a, D., Nanayakkara, O., Nityanandam, R., Mercado, N. B., Borducchi, E. N., Agarwal, A., Brinkman, A. L., Cabral, C., Chandrashekar, A., Giglio, P. B., Jetton, D., & Barouch, D. H. (2016). Protective efficacy of multiple vaccine platforms against Zika virus challenge in rhesus monkeys. Science, 353(6304), 1129–1132. 10.1126/science.aah6157

Ahi, Y. S., Bangari, D. S., & Mittal, S. K. (2011). Adenoviral Vector Immunity: Its implications and circumvention strategies. Current Gene Therapy, 11(4), 307-320. doi: 10.2174/156652311796150372.

Akello, J. O., Kamgang, R., Barbani, M. T., Suter-Riniker, F., Leib, S. L., & Ramette, A. (2020). Epidemiology of human adenoviruses: A 20-year retrospective observational study in hospitalized patients in Bern, Switzerland. Clinical Epidemiology, 12, 353–366. 10.2147/CLEP.S246352

Barnard, D. L., Wong, M.-H., Bailey, K., Day, C. W., Sidwell, R. W., Hickok, S. S., & Hall, T. J. (2007). Effect of oral gavage treatment with ZnAL42 and other metallo-ion formulations on influenza A H5N1 and H1N1 virus infections in mice. Antiviral Chemistry & Chemotherapy, 18(3), 125-132.

Belser, J. A., Katz, J. M., & Tumpey, T. M. (2011). The ferret as a model organism to study influenza A virus infection. Disease Models and Mechanisms, 4(5), 575–579. 10.1242/dmm.007823

Buckland, B. C. (2015). The development and manufacture of influenza vaccines. Human Vaccines and Immunotherapeutics, 11(6), 1357–1360. 10.1080/21645515.2015.1026497

Casimiro, D. R., Tang, A., Chen, L., Fu, T.-M., Evans, R. K., Davies, M.-E., Freed, D. C., Hurni, W., Aste-Amezaga, J. M., Guan, L., Long, R., Huang, L., Harris, V., Nawrocki, D. K., Mach, H., Troutman, R. D., Isopi, L. A., Murthy, K. K., Rice, K., & Shiver, J. W. (2003). Vaccine-induced immunity in baboons by using DNA and replication-incompetent adenovirus type 5 vectors expressing a human immunodeficiency virus type 1 gag gene. Journal of Virology, 77(13), 7663–7668. 10.1128/jvi.77.13.7663-7668.2003

Chen, C., Tang, T., Chen, Z., Chen, L., Cheng, J., Li, F., Sun, J., Zhao, J., Wang, Y., Yan, Q., Zhao, J., & Zhu, A. (2024). Antibody dynamics for heterologous boosters with aerosolized Ad5-nCoV following inactivated COVID-19 vaccines. Human Vaccines & Immunotherapeutics, 20(1), 2423466. 10.1080/21645515.2024.2423466

Cheshenko, N., Krougliak, N., Eisensmith, R. C., & Krougliak, V. A. (2001). A novel system for the production of fully deleted adenovirus vectors that does not require helper adenovirus. Gene Therapy, 8(11), 846–854. 10.1038/sj.gt.3301459

Coughlan, L., Sridhar, S., Payne, R., Edmans, M., Milicic, A., Venkatraman, N., Lugonja, B., Clifton, L., Qi, C., Folegatti, P. M., Lawrie, A. M., Roberts, R., de Graaf, H., Sukhtankar, P., Faust, S. N., Lewis, D. J. M., Lambe, T., Hill, A., & Gilbert, S. C. (2018). Heterologous two-dose vaccination with simian adenovirus and poxvirus vectors elicits long-lasting cellular immunity to influenza virus A in healthy adults. EBioMedicine, 29, 146–154. 10.1016/j.ebiom.2018.02.011

Crawford, K. H. D., Eguia, R., Dingens, A. S., Loes, A. N., Malone, K. D., Wolf, C. R., Chu, H. Y., Tortorici, M. A., Veesler, D., Murphy, M., Pettie, D., King, N. P., Balazs, A. B., & Bloom, J. D. (2020). Protocol and reagents for pseudotyping lentiviral particles with SARS-CoV-2 spike protein for neutralization assays. Viruses, 12(5), 513 10.3390/v12050513

Dormond, E., Chahal, P., Bernier, A., Tran, R., Perrier, M., & Kamen, A. (2010). An efficient process for the purification of helper-dependent adenoviral vector and removal of helper virus by iodixanol ultracentrifugation. Journal of Virological Methods, 165(1), 83–89. 10.1016/j.jviromet.2010.01.008

Dormond, E., Meneses-Acosta, A., Jacob, D., Durocher, Y., Gilbert, R., Perrier, M., & Kamen, A. (2009). An efficient and scalable process for helper-dependent adenoviral vector production using polyethylenimine-adenofection. Biotechnology and Bioengineering, 102(3), 800–810. 10.1002/bit.22113

Eglon, M. N., Duffy, A. M., O’Brien, T., & Strappe, P. M. (2009). Purification of adenoviral vectors by combined anion exchange and gel filtration chromatography. The Journal of Gene Medicine, 11(11), 978–989. 10.1002/jgm.1383

Fausther-Bovendo, H., & Kobinger, G. P. (2014). Pre-existing immunity against Ad vectors. Human Vaccines & Immunotherapeutics, 10(10), 2875–2884. 10.4161/hv.29594

Ferrara, F., & Temperton, N. (2018). Pseudotype neutralization assays: From laboratory bench to data analysis. Methods and Protocols, 1(1), 1–16. 10.3390/mps1010008

Gao, W., Soloff, A. C., Lu, X., Montecalvo, A., Nguyen, D. C., Matsuoka, Y., Robbins, P. D., Swayne, D. E., Donis, R. O., Katz, J. M., Barratt-Boyes, S. M., & Gambotto, A. (2006). Protection of mice and poultry from lethal H5N1 avian influenza virus through adenovirus-based immunization. Journal of Virology, 80(4), 1959–1964. 10.1128/jvi.80.4.1959-1964.2006

Graziosi, G., Lupini, C., Catelli, E., & Carnaccini, S. (2024). Highly pathogenic avian influenza (HPAI) H5 clade 2.3.4.4b virus infection in birds and mammals. Animals, 14(9), 1372. 10.3390/ani14091372

Happe, M., Hofstetter, A. R., Wang, J., Yamshchikov, G. V., Holman, L. S. A., Novik, L., Strom, L., Kiweewa, F., Wakabi, S., Millard, M., Kelley, C. F., Kabbani, S., Edupuganti, S., Beck, A., Kaltovich, F., Murray, T., Tsukerman, S., Carr, D., Ashman, C., & Matovu, R. (2024). Heterologous cAd3-Ebola and MVA-EbolaZ vaccines are safe and immunogenic in US and Uganda phase 1/1b trials. NPJ Vaccines, 9(1), 67. 10.1038/s41541-024-00833-z

He, J., & Kam, Y. W. (2024). Insights from avian influenza: a review of its multifaceted nature and future pandemic preparedness. Viruses, 16(3), 458. 10.3390/v16030458

Herfst, S., Schrauwen, E. J. A., Linster, M., Chutinimitkul, S., De Wit, E., Munster, V. J., Sorrell, E. M., Bestebroer, T. M., Burke, D. F., Smith, D. J., Rimmelzwaan, G. F., Osterhaus, A. D. M. E., & Fouchier, R. A. M. (2012). Airborne transmission of influenza A/H5N1 virus between ferrets. Science, 336(6088), 1534–1541. 10.1126/science.1213362

Imai, M., Watanabe, T., Hatta, M., Das, S. C., Ozawa, M., Shinya, K., Zhong, G., Hanson, A., Katsura, H., Watanabe, S., Li, C., Kawakami, E., Yamada, S., Kiso, M., Suzuki, Y., Maher, E. A., Neumann, G., & Kawaoka, Y. (2012). Experimental adaptation of an influenza H5 HA confers respiratory droplet transmission to a reassortant H5 HA/H1N1 virus in ferrets. Nature, 486(7403), 420–428. 10.1038/nature10831

Ison, M. G., & Marrazzo, J. (2025). The Emerging Threat of H5N1 to Human Health. New England Journal of Medicine, 392(9), 916–918. 10.1056/NEJMe2416323

Jia, S., Yin, Z., Pan, H., Wang, F., Liu, X., Wang, Q., Zhang, L., Tang, J., Yang, H., Du, J., Wang, Z., Jin, P., Peng, Z., Tang, R., Kang, G., Wang, X., Li, S., Wang, W., Li, J., & Zhu, F. (2024). Relative effectiveness of a heterologous booster dose with adenovirus type 5 vectored COVID-19 vaccine versus three doses of inactivated COVID-19 vaccine in adults during a nationwide outbreak of omicron predominance, in China: a retrospective, individually matched cohort-control study. Emerging Microbes and Infections, 13(1), 2332660. 10.1080/22221751.2024.2332660

Kamps, B. S., Hoffmann, C., & Preiser, W. (2006). Influenza Report 2016 (B. S. Kamps, C. Hoffmann, & W. Preissr, Eds.). Flying Publisher.

Kandeil, A., Patton, C., Jones, J. C., Jeevan, T., Harrington, W. N., Trifkovic, S., Seiler, J. P., Fabrizio, T., Woodard, K., Turner, J. C., Crumpton, J. C., Miller, L., Rubrum, A., DeBeauchamp, J., Russell, C. J., Govorkova, E. A., Vogel, P., Kim-Torchetti, M., Berhane, Y., & Webby, R. J. (2023). Rapid evolution of A(H5N1) influenza viruses after intercontinental spread to North America. Nature Communications, 14(1), 3082. 10.1038/s41467-023-38415-7

Kobinger, G. P., Figueredo, J. M., Rowe, T., Zhi, Y., Gao, G., Sanmiguel, J. C., Bell, P., Wivel, N. A., Zitzow, L. A., Flieder, D. B., Hogan, R. J., & Wilson, J. M. (2007). Adenovirus-based vaccine prevents pneumonia in ferrets challenged with the SARS coronavirus and stimulates robust immune responses in macaques. Vaccine, 25(28), 5220–5231. 10.1016/j.vaccine.2007.04.065

Koehler, D. R., Martin, B., Corey, M., Palmer, D., Ng, P., Tanswell, A. K., & Hu, J. (2006). Readministration of helper-dependent adenovirus to mouse lung. Gene Therapy, 13(9), 773–780. 10.1038/sj.gt.3302712

Krammer, F., Smith, G. J. D., Fouchier, R. A. M., Peiris, M., Kedzierska, K., Doherty, P. C., Palese, P., Shaw, M. L., Treanor, J., Webster, R. G., & García-Sastre, A. (2018). Influenza. Nature Reviews Disease Primers, 4(1), 3. 10.1038/s41572-018-0002-y

Lambert, L. C., & Fauci, A. S. (2010). Influenza vaccines for the future. New England Journal of Medicine. 21(18), 2036-2044. 10.1056/NEJMra1002842.

Lee, D., Liu, J., Junn, H. J., Lee, E. J., Jeong, K. S., & Seol, D. W. (2019). No more helper adenovirus: production of gutless adenovirus (GLAd) free of adenovirus and replication-competent adenovirus (RCA) contaminants. Experimental and Molecular Medicine, 51(10), 1-18. 10.1038/s12276-019-0334-z

Li, X., Liu, J., Li, W., Peng, Q., Li, M., Ying, Z., Zhang, Z., Liu, X., Wu, X., Zhao, D., Yang, L., Cao, S., Huang, Y., Shi, L., Xu, H., Wang, Y., Yue, G., Suo, Y., Nie, J., & Li, Y. (2023). Heterologous prime-boost immunisation with mRNA- and AdC68-based 2019-nCoV variant vaccines induces broad-spectrum immune responses in mice. Frontiers in Immunology, 14,1142394. 10.3389/fimmu.2023.1142394

Mellor, A. (1986). The class I MHC gene family in mice. Immunology Today. 7(1),19–24.

Morse, M. A., Chaudhry, A., Gabitzsch, E. S., Hobeika, A. C., Osada, T., Clay, T. M., Amalfitano, A., Burnett, B. K., Devi, G. R., Hsu, D. S., Xu, Y., Balcaitis, S., Dua, R., Nguyen, S., Balint, J. P., Jones, F. R., & Lyerly, H. K. (2013). Novel adenoviral vector induces T-cell responses despite anti-adenoviral neutralizing antibodies in colorectal cancer patients. Cancer Immunology, Immunotherapy, 62(8), 1293–1301. 10.1007/s00262-013-1400-3

Olsen, B., Munster, V. J., Wallensten, A., Waldenström, J., Osterhaus, A. D. M. E., & Fouchier, R. A. M. (2006). Global patterns of influenza A virus in wild birds. Science, 312(5772), 384–388. 10.1126/science.1122438

Palmer, D. J., & NG, P. (2005). Helper-dependent adenoviral vectors for gene therapy. Human Gene Therapy, 16(1), 1–16. 10.1089/hum.2005.16.1

Palmer, D., & Ng, P. (2003). Improved system for helper-dependent adenoviral vector production. Molecular Therapy, 8(5), 846–852. 10.1016/j.ymthe.2003.08.014

Parrish, C. R., Murcia, P. R., & Holmes, E. C. (2015). Influenza virus reservoirs and intermediate hosts: dogs, horses, and new possibilities for influenza virus exposure of humans. Journal of Virology, 89(6), 2990–2994. 10.1128/jvi.03146-14

Patel, A., Tikoo, S., & Kobinger, G. (2010). A porcine adenovirus with low human seroprevalence is a promising alternative vaccine vector to human adenovirus 5 in an H5N1 virus disease model. PLoS ONE, 5(12), e15301. 10.1371/journal.pone.0015301

Qi, Y., Tarbet, E. B., Cull, J. W., Zhang, X., & Staerz, U. D. (2024). A Potent Pandemic Avian influenza virus vaccine based on a 4th generation fully deleted adenoviral vector. BioRxiv. 10.1101/2024.12.30.630761

Qi, Y., Zhang, X., Wheeler Cull, J., Wall, C., Maslanik, W., & D. Staerz, U. (2024). The Flexible, Versatile and Fast GreGT Platform of Fully Deleted Helper-Virus Independent Adenoviral Vectors. Archives of Microbiology & Immunology, 8(4). 493-502. 10.26502/ami.936500201

Refaey, S., Azziz-Baumgartner, E., Amin, M. M., Fahim, M., Roguski, K., Elaziz, H. A. E. A., Iuliano, A. D., Salah, N., Uyeki, T. M., Lindstrom, S., Davis, C. T., Eid, A., Genedy, M., & Kandeel, A. (2015). Increased number of human cases of influenza virus A(H5N1) infection, Egypt, 2014–15. Emerging Infectious Diseases, 21(12), 2171–2173. 10.3201/eid2112.150885

Romero, P., Corradinj, G., Luescher, I. F., & Maryanski, J. L. (1991). H-2Kd-restricted antigenic peptides share a simple binding motif. Journal of Experimental Medicine, 174, 603-612.

Roy, S., Kobinger, G. P., Lin, J., Figueredo, J., Calcedo, R., Kobasa, D., & Wilson, J. M. (2007). Partial protection against H5N1 influenza in mice with a single dose of a chimpanzee adenovirus vector expressing nucleoprotein. Vaccine, 25(39–40), 6845–6851. 10.1016/j.vaccine.2007.07.035

Russell, W. C., Graham, F. L., Smiley, J., & Nairn, R. (1977). Characteristics of a human cell line transformed by DNA from human adenovirus type 5. Journal of General Virology, 36(1), 59–72. 10.1099/0022-1317-36-1-59

Singh, N., Pandey, A., Jayashankar, L., & Mittal, S. K. (2008). Bovine adenoviral vector-based H5N1 influenza vaccine overcomes exceptionally high levels of pre-existing immunity against human adenovirus. Molecular Therapy, 16(5), 965–971. 10.1038/mt.2008.12

Sutton, T. C. (2018). The pandemic threat of emerging H5 and H7 avian influenza viruses. Viruses 10(9), 461. 10.3390/v10090461

Tapia, M. D., Sow, S. O., Lyke, K. E., Haidara, F. C., Diallo, F., Doumbia, M., Traore, A., Coulibaly, F., Kodio, M., Onwuchekwa, U., Sztein, M. B., Wahid, R., Campbell, J. D., Kieny, M.-P., Moorthy, V., Imoukhuede, E. B., Rampling, T., Roman, F., De Ryck, I., & Levine, M. M. (2016). Use of ChAd3-EBO-Z Ebola virus vaccine in Malian and US adults, and boosting of Malian adults with MVA-BN-Filo: a phase 1, single-blind, randomised trial, a phase 1b, open-label and double-blind, dose-escalation trial, and a nested, randomised, double-blind, placebo-controlled trial. The Lancet Infectious Diseases, 16(1), 31–42. 10.1016/S1473-3099(15)00362-X

Temperton, N. J., Hoschler, K., Major, D., Nicolson, C., Manvell, R., Hien, V. M., Ha, D. Q., de Jong, M., Zambon, M., Takeuchi, Y., & Weiss, R. A. (2007). A sensitive retroviral pseudotype assay for influenza H5N1-neutralizing antibodies. Influenza and Other Respiratory Viruses, 1(3), 105–112. 10.1111/j.1750-2659.2007.00016.x

Uyeki, T., Milton, S., Webb, C., Shetty, V., Kniss, K., Frederic, J., De la Cruz, J., Lidell, J., Di, H., Kirby, M., Barnes, J., & Davis, C. (2025). Highly Pathogenic Avian Influenza A(H5N1) Virus Infection in a Dairy Farm Worker. New England Journal of Medicine, 390(21), 2028–2019.

Venkatesh, D., Poen, M. J., Bestebroer, T. M., Scheuer, R. D., Vuong, O., Chkhaidze, M., Machablishvili, A., Mamuchadze, J., Ninua, L., Fedorova, N. B., Halpin, R. A., Lin, X., Ransier, A., Stockwell, T. B., Wentworth, D. E., Kriti, D., Dutta, J., van Bakel, H., Puranik, A., & Lewis, N. S. (2018). Avian influenza viruses in wild birds: virus evolution in a multihost ecosystem. Journal of Virology, 92(1), e00433-18. 10.1128/jvi.00433-18

Weaver, E. A., Nehete, P. N., Buchl, S. S., Senac, J. S., Palmer, D., Ng, P., Sastry, K. J., & Barry, M. A. (2009). Comparison of replication-competent, first generation, and helper-dependent adenoviral vaccines. PLoS ONE, 4(3), e5059 10.1371/journal.pone.0005059

Weaver, E. A., Nehete, P. N., Nehete, B. P., Yang, G., Buchl, S. J., Hanley, P. W., Palmer, D., Montefiori, D. C., Ferrari, G., Ng, P., Sastry, K. J., & Barry, M. A. (2013). Comparison of Systemic and Mucosal Immunization with Helper-Dependent Adenoviruses for Vaccination against Mucosal Challenge with SHIV. PLoS ONE, 8(7)., e67574,. 10.1371/journal.pone.0067574

Webby, R., & Uyeki, T. (2024). An Update on Highly Pathogenic Avian Influenza A(H5N1) Virus, Clade 2.3.4.4b. Journal of Infectious Diseases, 23-(3), 533–542.

